# Hierarchical neuronal processing in primary somatosensory cortex during action observation

**DOI:** 10.64898/2026.07.01.735792

**Authors:** Tomomichi Oya, Amit Yaron, Joachim Confais, Shinji Kubota, Satomi Kikuta, Kazuhiko Seki

## Abstract

Voluntary movement requires the central nervous system to transform and integrate visual and somatosensory information into coordinated motor outputs. Although mirror-like neuronal activity during both action execution and observation has been extensively described in premotor, motor, and parietal cortices, it remains unknown whether the primary somatosensory cortex (S1) also participates in the action observation network. Here, we recorded single-unit activity from cytoarchitectonically defined areas 3a, 3b, 1, and 2 in macaque S1 while monkeys either executed or observed grasping movements. Approximately one-third of neurons across S1 modulated their firing during action observation, with the proportion of responsive neurons increasing from area 3 to areas 1 and 2, consistent with the hierarchical organization of somatosensory processing. Most action observation neurons showed congruent activity during action execution and observation, suggesting that these responses may reflect top-down motor-related or integrated visuomotor signals and are unlikely to be explained by visual input alone. The higher prevalence of action observation neurons in areas 1 and 2 suggests that action observation-related signals preferentially influence later stages of somatosensory processing, potentially via cortico-cortical interactions with motor and parietal regions.

## 1 Introduction

The central nervous system is capable of selectively processing and integrating various sensory inputs to generate appropriate motor outputs—functions that are critical to voluntary movement. Among these sensory modalities, vision and somatosensation play central roles in the dexterous and skilled manual grasping in primates (Mundinano et al. (2018); Davare et al. (2011)). Traditionally, visuomotor transformation and somatosensory-to-motor transformation have often been described as separate processes: visuomotor transformation converts visual information about object location and features into movement commands (e.g., Graziano et al. (1994)), whereas somatosensory-to-motor transformation uses tactile and proprioceptive feedback to dynamically shape motor output through transcortical sensorimotor pathways (Evarts (1973)). However, accumulating evidence suggests that these modalities are tightly interdependent rather than functionally isolated. The neurons in the premotor and parietal cortices can represent visual space in body- or limb-centered coordinates (Graziano et al. (1994)), and the premotor neurons integrate visual information about the position of the limb with proprioceptive signals from the arm itself (Graziano et al. (2000)). Moreover, sensorimotor areas frequently exhibit mixed representations of visual, somatosensory, and motor-related signals during reaching and grasping behaviors (Shen and Alexander (1997); Murata et al. (2000)). These findings suggest that visuomotor and somatosensory transformations continuously interact and may rely on partially overlapping neural substrates. However, how and where these transformation pathways converge and undergo integration in the brain is still incompletely understood. Clarifying this integration is essential to understanding how goal-directed movements such as reaching and grasping are organized.

A key insight into visuomotor integration emerged from the discovery of mirror neurons—neurons that fire both when an individual performs an action and when observing another perform the same action. First identified in the macaque premotor cortex (di Pellegrino et al. (1992); Gallese et al. (1996); Rizzolatti et al. (1996), Ferrari et al. (2005), Bonini et al. (2014), Caggiano et al. (2011), Caggiano et al. (2013), Caggiano et al. (2015)), such neurons have also been found in the parietal cortex (Taira et al. (1990); Sakata et al. (1995); Gallese et al. (2002); Fogassi et al. (2005); Fujii et al. (2007)) and supplementary motor area (Mukamel et al. (2010)). These findings implicate a fronto-parietal network—linking the visual dorsal stream to the parietal and frontal lobes—as a substrate for visuomotor transformation.

Given the functional properties of these higher-order areas, activity during action observation has been interpreted in two main ways. One interpretation posits that such activity reflects a cognitive process aimed at understanding others’ actions, intentions, or emotional states within a social context (Rizzolatti and Craighero (2004); Iacoboni et al. (2005); Caggiano et al. (2012); Keysers and Gazzola (2009); Rizzolatti and Sinigaglia (2016)). An alternative view holds that the observer internally simulates the observed action, thereby refining their own motor representations or facilitating motor learning (Jeannerod (2001); Brass and Heyes (2005); Mattar and Gribble (2005)). These hypotheses are not mutually exclusive; however, the former emphasizes cognitive understanding, whereas the latter engages mechanisms of motor execution and internal modeling.

Recent studies have extended mirror neuron findings into primary motor cortex (M1) (Tkach et al. (2007); Dushanova and Donoghue (2010); Mazurek et al. (2018); Jerjian et al. (2020)), including corticospinal neurons, i.e., the output neurons of the motor system (Kraskov et al. (2009, 2014)). This supports the simulation hypothesis, which proposes that observing an action triggers internal rehearsal, including the sensory consequences of movement (Jeannerod (2001); Keysers et al. (2010)). If so, somatosensory regions should also be engaged during action observation.

Indeed, studies in humans have shown that S1 activity can increase during the observation or imagery of movement, even in the absence of direct peripheral stimulation (Jafari et al. (2020)), and that visual stimuli aligned with tactile input can activate somatotopic maps in S1 (Kuehn et al. (2018)). These findings suggest that the neurons in S1 – the first cortical area for somatosensory processing – could also exhibit mirror-like activity. Demonstrating such activity in S1 would provide important evidence that somatosensory cortex participates in neural processes engaged during action observation.

Given known anatomical connections from premotor, motor and parietal areas to S1 (Gharbawie et al. (2011); Yazdan-Shahmorad et al. (2018); Padberg et al. (2019)), it is plausible that top-down motor-related signals reach early somatosensory cortex during action observation. However, no systematic studies to date have investigated single-neuron activity during action observation across the subfields of S1, nor is it known at which level of its hierarchical processing within S1 such signals are expressed.

To address this, we recorded and analyzed neuronal activity from precisely defined areas 3a, 3b, 1, and 2 in S1 of macaques during both self-executed and observed grasping movements. Our findings revealed that a sizable proportion (~30%) of neurons in S1 responded during action observation. Among these, area 3a and 3b showed the smallest proportion of responsive neurons, while areas 1 and 2 exhibited a greater proportion. This pattern parallels the hierarchical progression of sensory processing from lower to higher-order regions within S1. Moreover, neurons in area 2 showed greater incongruence between execution and observation conditions compared with other subdivisions, suggesting that action observation-related activity in area 2 may reflect inputs or processing distinct from those in other S1 areas. These results suggest that visuomotor information related to action observation is distributed non-uniformly across the hierarchical organization of S1, putatively paralleled with hierarchical integration, along which receptive fields expand and modalities converge.

## 2 Results

### 2.1 Neurons in the primary somatosensory cortex were modulated during action observation as well as voluntary task execution

We asked whether S1 contained action observation neurons – neurons showing significant modulation during both voluntary grasp execution and action observation. To this end, we trained monkeys to grasp two different objects and to observe the same grasping behaviors performed by the human experimenter without overt movements (Figure 1A, B). The objects presented were the ring and precision grip objects for monkey Mu, and the ball and precision grip objects for monkey Ab; the selection was tailored to each animal’s preference. To characterize the behavior that could modulate the activity of S1 neurons during grasp execution and observation, the task sequence was segmented into 8 epochs with respect to behavioral events: ‘Object Presentation’, ‘Go Cue’, ‘Reach Onset’, ‘Middle of Reach’, ‘Item Touch’, ‘Item Hold Start’, ‘Release Cue’, and ‘Item Hold End’ (Figure 1C; for detailed definitions of the epochs, see the Methods).

**Fig. 1.**
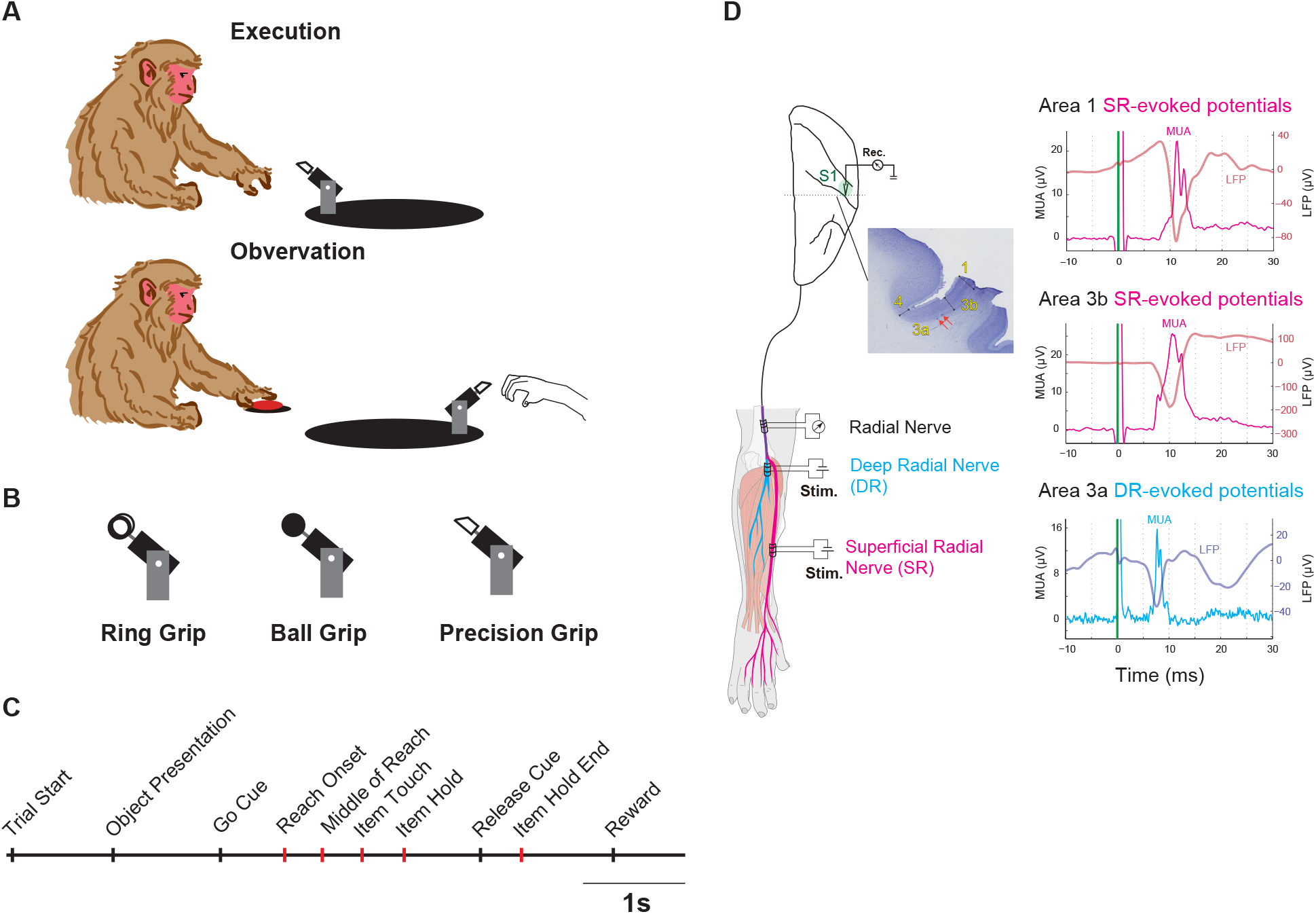
**A**. Behavioral Task. In the Execution condition, the monkey performed a grip task, whereas in Observation condition, the monkey watched the human experimenter perform the same grip while laying its hand on the home button. **B**. Grip objects. Two of three grip objects (Ring, Ball, or Precision grip) were presented to each monkey **C**. Task sequence. Black ticks are the events defined by the task scheduler, and red ticks are the events defined based on the animal’s behavior. **Trial Start** - The trial began when both the monkey and the human experimenter placed one hand on the homepad for a certain period of time (100 ms). **Object Presentation** - After a delay period (1000 ms), the grip object was presented to the monkey and turned visible. **Go Cue** - After another period of delay (1000 ms), another color of LED illumination and a beep sound were given, which instructed the monkey to reach out, grasp and pull the object. **Reach Onset** - the time the monkey released the homepad. **Middle of Reach** - the time in the middle of Reach Onset and the time the monkey touched the object. **Item Touch** - the time when a custom-made electrostatic sensor is activated. **Item Hold** - the time when the monkey pulled the object to the 90% of range of motion. **Release Cue** - the time when the beep sound was given to signal the monkey to release the object. **Item Hold End** - the time the lever of the object returned below 10% of range of motion. **Reward** - the beep sound and juice or food reward were given. **D**. Recording and identification of primary somatosensory cortex and peripheral stimulation. Electrophysiological recording was made in the primary somatosensory cortex (S1). The recording sites for Brodmann’s areas inside S1 - areas 1, 3b, and 3a were classified based on receptive field, post-hoc anatomical examination, and the somatosensory evoked potentials (SEPs) elicited by stimulation of peripheral nerves carrying different modalities, i.e., superficial branch of radial nerve (SR) for a cutaneous afferent and deep branch of radial nerve (DR) for a muscle afferent. Areas 1 and 3b were determined by the responsiveness to SR, and further classified by the shape and latency of the evoked potential, i.e., earlier response without positive deflection for area 3b versus late response with positive deflection for area 1. Area 3a exclusively responded to DR stimulation as it preferentially receives the proprioceptive input from the periphery.

To identify the subdivisions within S1, i.e., 3a, 3b, 1 and 2, we performed mapping sessions under sedation using electrical and tactile stimulation. Using coregistered structural images obtained by MRI and CT (Miocinovic et al. (2007)), we identified S1 in the postcentral gyrus (Supplemental Figure S1).

Within the digit representation of S1, we explored neurons and examined their responses by refining the technique we developed previously (Yamada et al. (2016)); using the combination of tactile palpation of the forearm and hand and electrical stimulation of the superficial radial (SR) and deep radial (DR) nerve branches, we were able to delineate the subdivisions more robustly and extensively. Via SR branch stimulation, we elicited the largest somatosensory evoked potential (SEP) in the thumb- and index finger-innervation zones in areas 3b and 1, while we evoked SEPs exclusively in area 3a across digits via DR branch stimulation (Figure 1D). Because SR contains fibres originating from cutaneous receptors, SR branch stimulation elicits SEPs in areas that receive cutaneous inputs, e.g., areas 3b and 1. In contrast, DR branch contains fibres from proprioceptive receptors, and its stimulation leads to SEPs in areas that receive proprioceptive inputs, e.g., area 3a. Based on the distinct presence of SEPs via different modalities, areas 3a and 3b were dissociated by the presence or absence of SEPs evoked by SR or DR at the anterior bank of the postcentral gyrus (Figure 1D). Areas 3b and 1 were also delineated based on a combination of SEP shapes elicited by SR stimulation (Figure 1D) and receptive fields; for area 1, a positive deflection preceded the negative deflection in the SEP, whereas no such positive deflection was observed for area 3b. This distinction in the SEP corresponded to neuronal characteristics that changed across the 1/3b border, including a reversal of the proximal-distal somatotopic gradient and a broadening of the digit receptive fields (Goodman et al. (2019)).

Areas 1 and 2 were delineated by their rostro-caudal position along the cortical surface and the complexity of the receptive fields, as well as responsiveness to the mixed modalities of electrical stimulation, i.e., a mosaic-like patch of responsiveness to either or both SR and DR stimulation. In addition to these separate sessions for mapping the subdivisions, we routinely stimulated the peripheral nerves for confirmation before recording during behavioral sessions. The delineation was finally confirmed or adapted using histological examination of cortical cytoarchitecture after the animal was sacrificed (Figure 1D and Supplementary Figure S2), with reference to previous literature (Nelson et al. (1980) and Goodman et al. (2019)).

From these identified subdivisions in S1, we recorded a total of 374 neurons during action execution and observation from two macaque monkeys (195 neurons from monkey Mu and 179 neurons from monkey Ab). After applying the trial-number criterion described in the Methods, all of these neurons were retained for analysis. We then restricted subsequent analyses to neurons that exhibited significant modulation in at least one epoch during execution of the grasping task relative to baseline activity, resulting in 190 neurons from monkey Mu and 170 neurons from monkey Ab.

An example neuron is shown in Figures 2-5. This neuron was classified as an area 3a neuron based on the criteria described above. It showed increased firing during action execution, peaking at the ‘Reach Onset’ epoch with additional activity at the ‘Item Hold End’ epoch, for both Ring (Figure 2) and Precision Grip (Figure 3) conditions. This firing characteristic was largely consistent with that of area 3a neurons reported in the previous literature Goodman et al. (2019). Notably, this neuron was also observation-active: its firing rate increased in the ‘Item Hold End’ epoch while the monkey observed the experimenter grasping both Ring (Figure 4) and Precision Grip (Figure 5) objects.

**Fig. 2.**
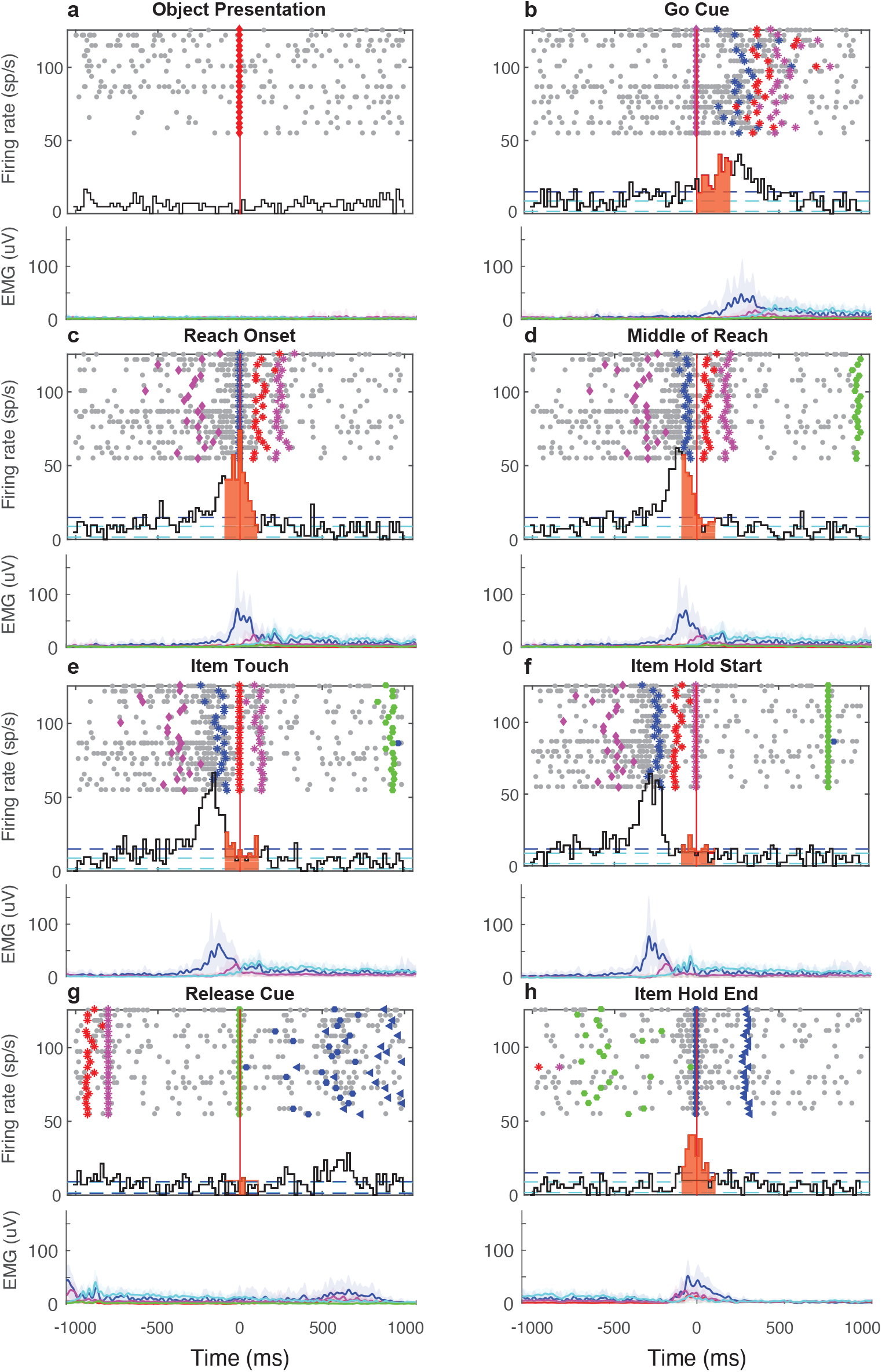
An exemplar S1 neuron in area 3a that showed modulation during both action execution and observation for different grasping objects. **a-h**. Upper panel: peri-event time histogram (PETH) of neuronal modulation; lower panel: electromyography (EMG) of hand muscles, aligned to each defined event: **a - Object Presentation, b - Go Cue, c - Reach Onset, d - Middle of Reach, e - Item Touch, f - Item Hold Start, g - Release Cue**, and **h - Item Hold End**. In the upper panel, dashed lines indicate the mean and 2SD, and significant epochs are color-shaded. In the lower panel, each EMG trace (mean and shaded 2SD) represents Extensor Digitorum Communis (EDC) - blue, Extensor Carpi Ulnaris (ECU) - red, Flexor Carpi Radialis (FCR) - magenta, Flexor Digitorum Superficialis (FDS) - cyan, and Triceps Brachii - green. Action execution during grasping of the ring object.

**Fig. 3.**
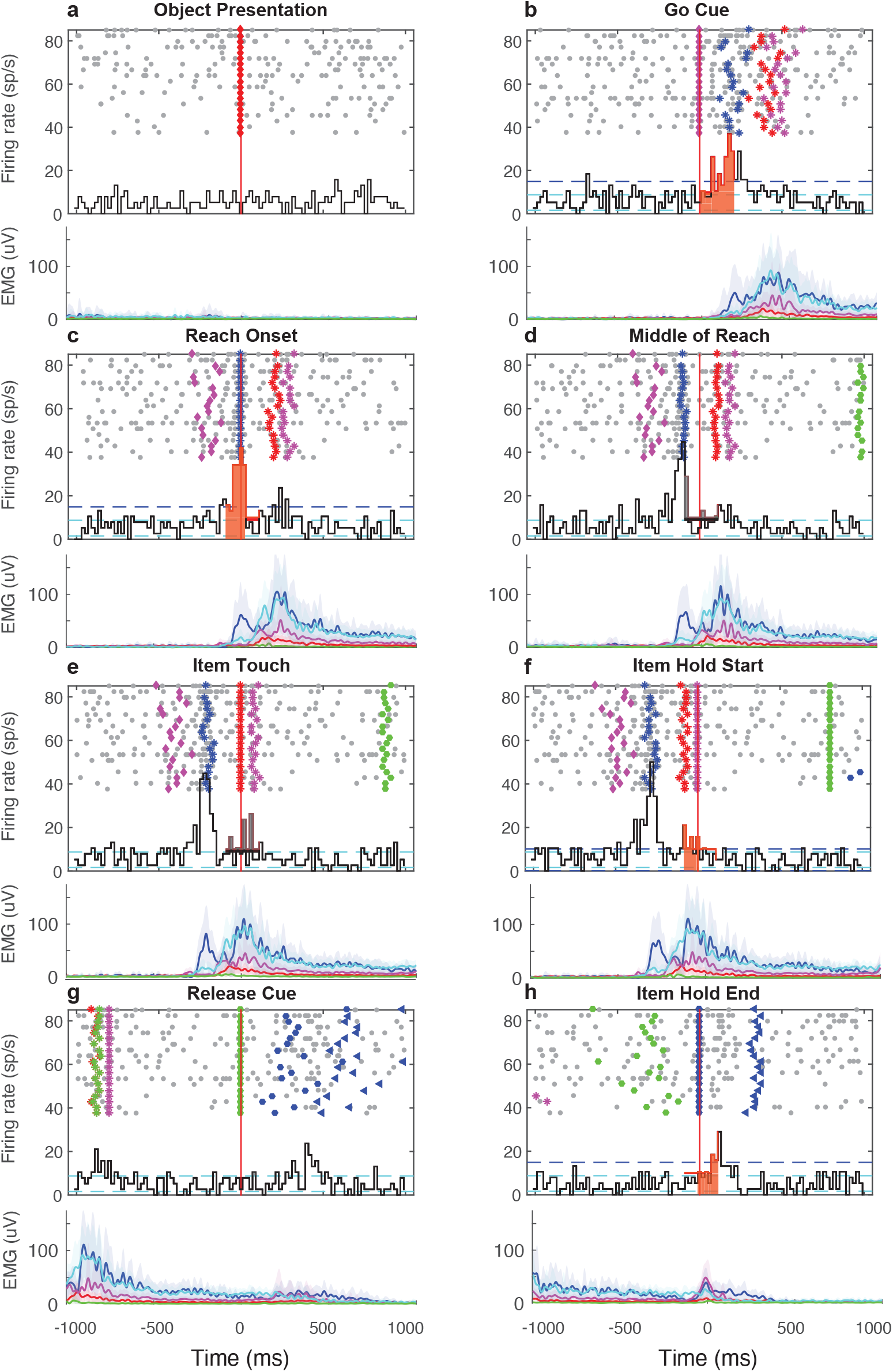
Action execution during grasping of the precision grip object. Cf. Figure 2 for detailed description of symbols and colors.

**Fig. 4.**
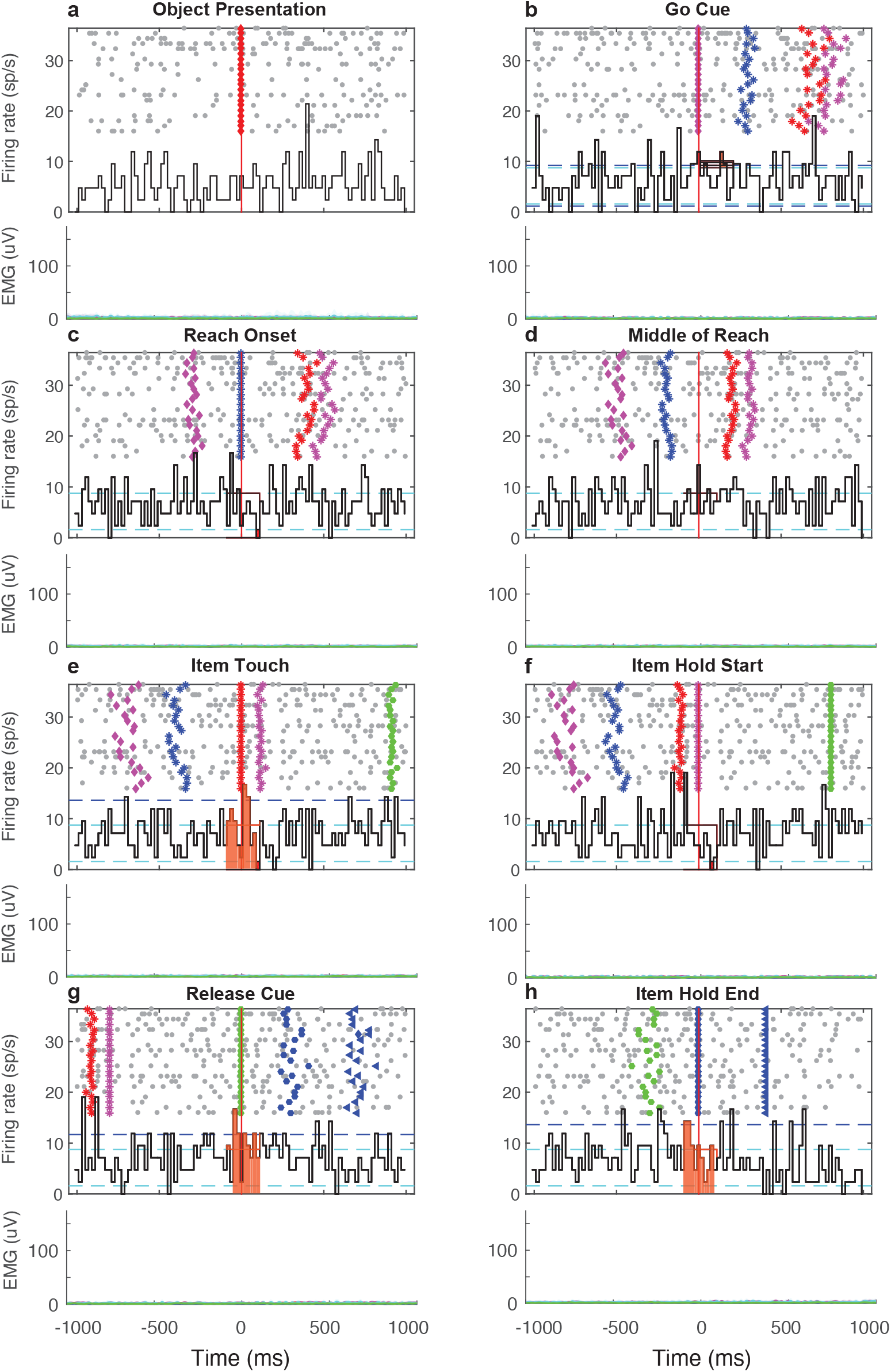
Action observation during grasping of the Ring object. Cf. Figure 2 for detailed description of symbols and colors.

**Fig. 5.**
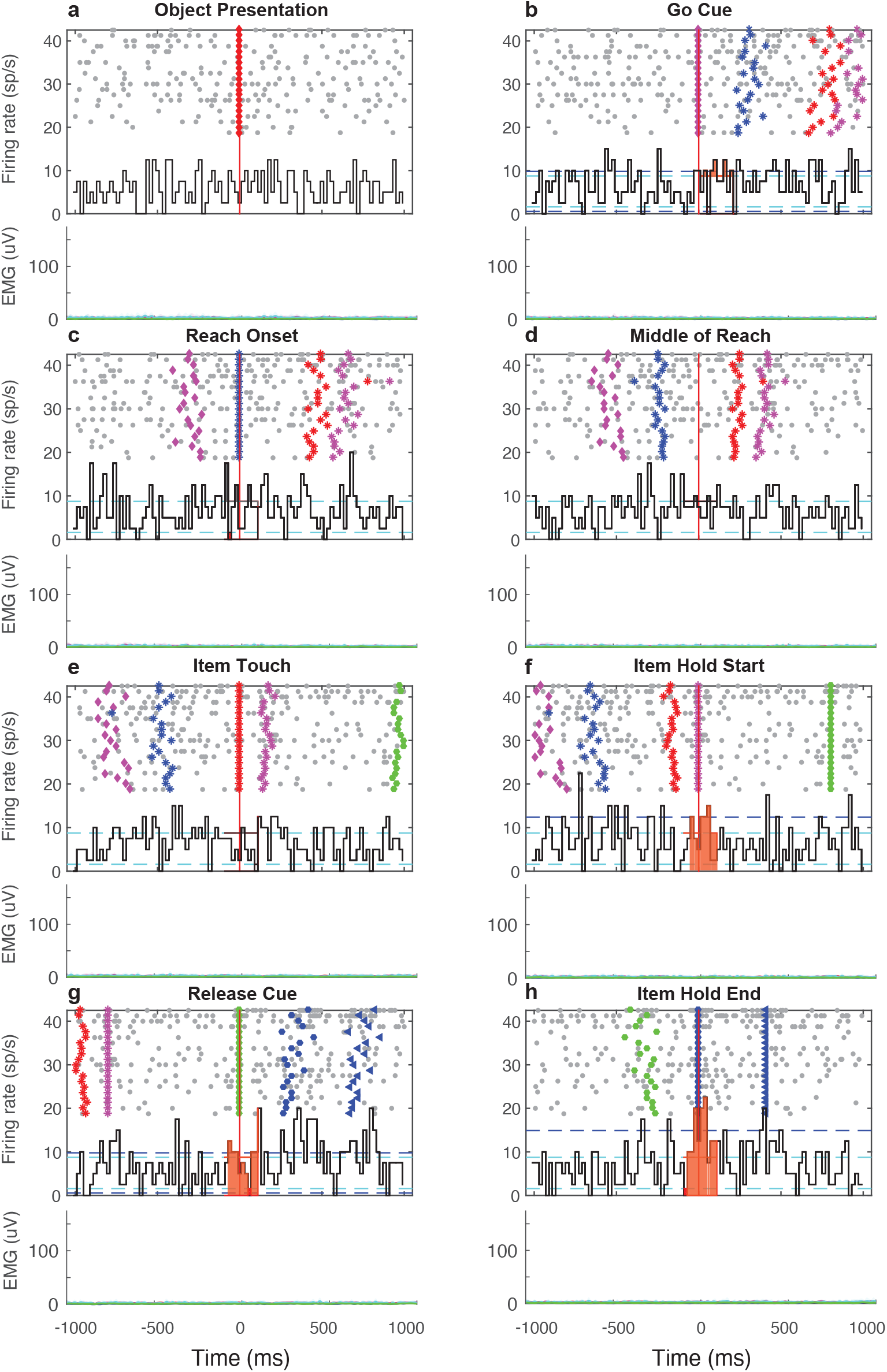
Action observation during grasping of the precision grip object. Cf. Figure 2 for detailed description of symbols and colors.

To rule out covert movements or muscle contractions as a source of observation-related activity, we recorded electromyography (EMG) from the forearm and upper arm muscles (Figures 2-5; blue: Extensor Digitorum Communis (EDC), red: Extensor Carpi Ulnaris (ECU), magenta: Flexor Carpi Radialis (FCR), cyan: Flexor Digitorum Superficialis (FDS), and green: Triceps Brachii (Triceps)), and confirmed that there was no significant increase in EMG activity beyond the baseline epoch (‘Object Presentation’) during the epochs in which the monkey observed the grasping behaviors, as shown in the EMG traces in the lower panels of Figure 4 and Figure 5. Among neurons that were activated during execution of the grasping tasks, approximately one-third showed an increase or decrease in activity in at least one behaviorally relevant epoch during observation of the grasping tasks in both monkeys (30.0% for monkey Mu and 35.2% for monkey Ab; Figure 6A). These neurons showed more significant epochs during action execution than during action observation (monkey Mu: Ring Execution, 3.62 ± 2.21; Precision Grip Execution, 4.14 ± 1.94; Ring Observation, 1.35 ± 1.69; Precision Grip Observation, 1.54 ± 1.40; monkey Ab: Ball Execution, 4.02 ± 1.91; Precision Grip Execution, 3.60 ± 1.97; Ball Observation, 2.03 ± 2.28; Precision Grip Observation, 1.68 ± 2.16; Figure 6B). We found no neurons in which the number of significant epochs was greater during action observation than during action execution.

**Fig. 6.**
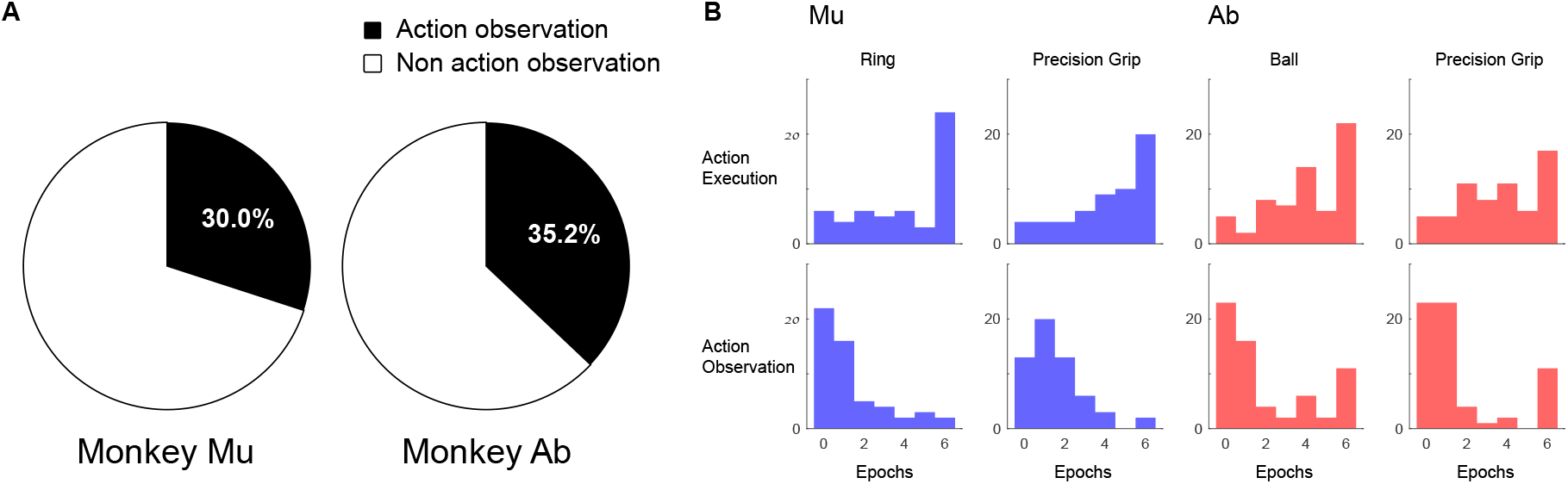
Population of action observation neurons in S1. A. Proportions of action observation neurons versus non-action observation neurons for each monkey (Mu and Ab). B. Histograms for the number of significant epochs during action execution and action observation. For monkey Mu, the ring and precision grip objects were grasped, whereas for monkey Ab, the ball-shaped and precision grip objects were used.

### 2.2 The proportion of neurons that were modulated during action observation was higher in areas 1 and 2, and lower in areas 3a/3b

We next compared the proportions of action observation neurons among different Brodmann areas in S1, i.e., areas 3a, 3b, 1 and 2. As shown in Figure 7, for each monkey, action observation neurons were less prevalent in areas 3a and 3b (Mu: 20.0% and 22.0%; Ab: 36.6% and 27.8%) than in areas 1 (Mu: 52.0%; Ab: 42.0%) and 2 (Mu: 64.7%; Ab: 34.1%). To statistically assess the hierarchical gradient, we pooled neurons into two groups based on their hierarchical position in S1: a lower-order group (areas 3a and 3b) and a higher-order group (areas 1 and 2). The proportion of action observation-responsive neurons was significantly higher in the higher-order group than in the lower-order group (Fisher’s exact test: 58/213 vs. 62/131, p *<* 0.001). This group-level difference supports a hierarchical gradient of action observation-related neuronal recruitment across S1 subdivisions.

**Fig. 7.**
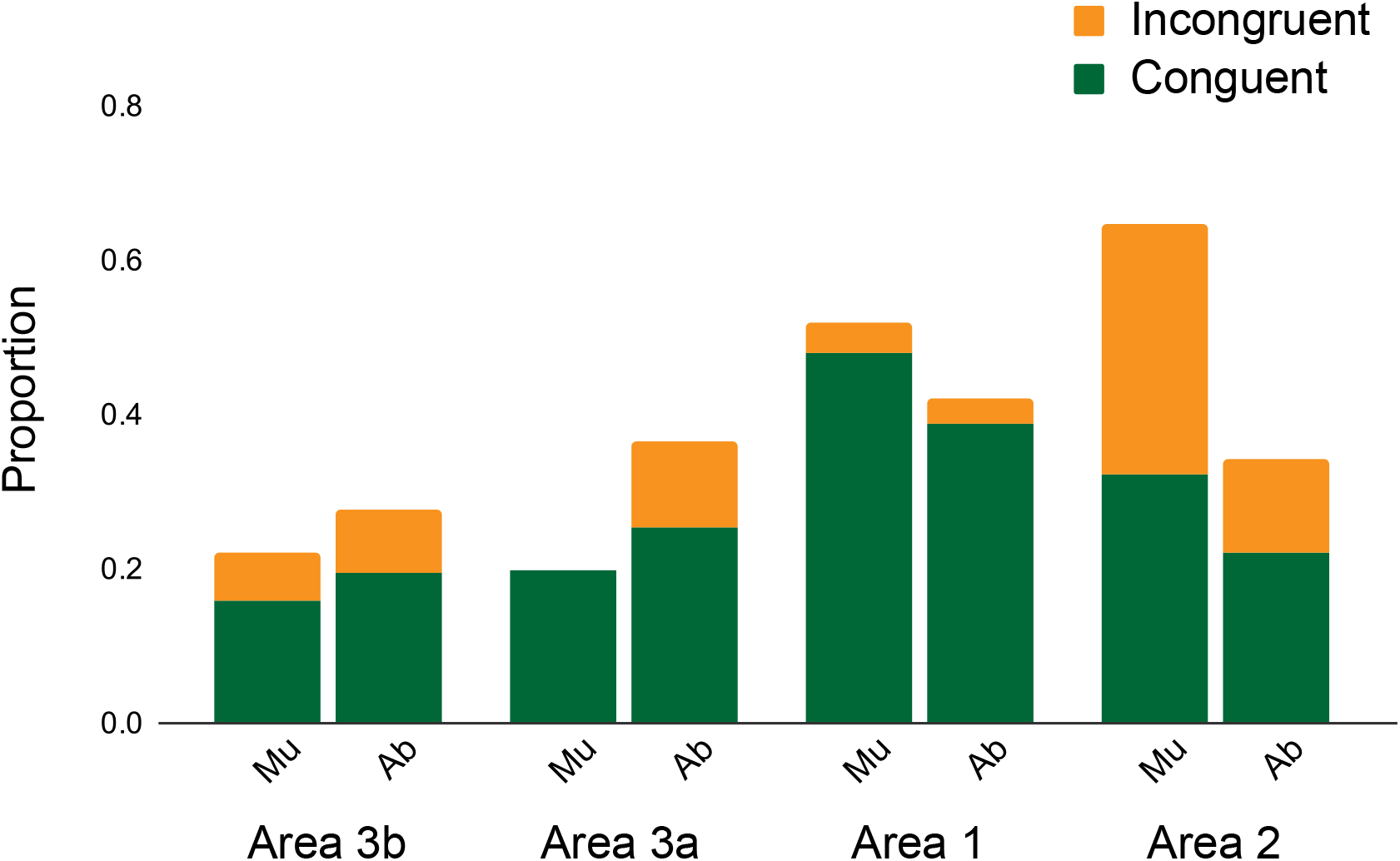
Proportions of action observation neurons and congruent neurons across subdivisions of S1. Green bars indicate neurons showing congruent activation between action execution and observation for the same grip type, whereas orange bars indicate incongruent neurons. The proportion of action observation-responsive neurons was significantly higher in the higher-order group (areas 1 and 2) than in the lower-order group (areas 3a and 3b) (Fisher’s exact test: 62/131 vs. 58/213, p *<* 0.001)

### 2.3 The epochs in which neurons were activated during action observation were congruent with the epochs in which neurons were significant during action execution

As originally defined (Rizzolatti and Craighero (2004)), mirror neurons exhibit specificity for both object and behavioral epoch, resulting in congruent activity during action execution and observation. To test whether the action observation neurons in our dataset showed similar specificity, we examined whether each neuron discharged for the same object and in the same epochs during observation as during execution. Overall, the majority of action observation neurons showed congruent activation for both epoch and object between action execution and observation (73.7% for monkey Mu, 73.0% for monkey Ab; green in Figure 7). Among S1 subdivisions, action observation neurons in area 1 showed the largest proportions of congruent neurons (92.3% for both monkeys; green bars in area 1 in Figure 7), whereas action observation neurons in area 2 showed the largest proportions of incongruent neurons in both monkeys (50.0% for monkey Mu and 64.3% for monkey Ab; orange bars in area 2 in Figure 7).

## 3 Discussion

In the present study, we aimed to investigate neuronal responses during action observation in S1. We found neurons that modulated their firing rate during both action execution and action observation across subdivisions in S1, specifically areas 3a, 3b, 1, and 2. While it is well established that neurons in S1 modulate their firing during movement because various cutaneous and proprioceptive receptors are activated during movement execution (Prud’homme and Kalaska (1994); Goodman et al. (2019)), neuronal responses during action observation have not previously been described in S1 at the single-cell level. Thus far, neurons exhibiting action observation-related activity have mainly been reported in motor-related cortical regions, including premotor cortex (di Pellegrino et al. (1992); Gallese et al. (1996); Rizzolatti et al. (1996); Ferrari et al. (2005); Bonini et al. (2014); Caggiano et al. (2011, 2013, 2015)), supplementary motor area (Mukamel et al. (2010)), primary motor cortex (Tkach et al. (2007); Dushanova and Donoghue (2010); Kraskov et al. (2014); Mazurek et al. (2018); Jerjian et al. (2020)), and parietal cortex (Taira et al. (1990); Sakata et al. (1995); Murata et al. (2000); Gallese et al. (2002); Fogassi et al. (2005); Fujii et al. (2007)). These regions are considered key nodes of fronto-parietal sensorimotor networks involved in visuomotor transformation and motor control. Although mirror-like activity in S1 has previously been suggested in human neuroimaging studies (Gazzola and Keysers (2009)), the present study provides direct neurophysiological evidence of such neuronal activity at the single-cell level across all four subdivisions of S1. The findings suggest that neuronal activity in S1 is not solely driven by somatosensory afferent signals ascending from the periphery, but may also reflect signals related to observing the actions of others. Such signals could arise from visual, motor-related, and/or integrated sensorimotor processes. In this sense, S1 may participate more broadly in distributed sensorimotor networks engaged during action observation. At the same time, the precise origin and functional significance of these responses remain unclear. These responses may arise through associative visuomotor interactions or top-down influences from motor-related cortical areas. Either way, the presence of action observation-related activity in S1 suggests that somatosensory cortex may participate in neural processes linking visuomotor and somatosensory-motor transformations, supporting interactions between these traditionally distinct sensorimotor pathways.

In the following sections, we discuss these neuronal responses in terms of the types of information they may represent – visually driven, motor-related, or integrated sensorimotor signals. We then consider the possible implications of these responses for action observation and voluntary movement control.

### 3.1 Origin and nature of action observation-related signals in S1

Since the seminal discovery of neurons modulated during action observation in premotor cortex (di Pellegrino et al. (1992)), accumulating evidence has demonstrated action observation-related activity across premotor, supplementary motor, primary motor, and parietal cortices. These findings suggest that signals related to action observation are broadly distributed throughout the fronto-parietal sensorimotor network involved in visuomotor transformation and motor control (Jeannerod et al. (1995)). The present study extends these observations by demonstrating action observation-related neuronal activity throughout subdivisions of S1. The action observation-related responses observed in S1 were generally weaker than those observed during action execution, suggesting that S1 is unlikely to be a primary source of such activity. Rather, the responses may reflect cortico-cortical influences from motor- and visuomotor-related cortical regions engaged during action observation. Dense reciprocal connections between M1 and S1 (Gharbawie et al. (2011); Yazdan-Shahmorad et al. (2018)), as well as known inputs from parietal association areas, provide plausible anatomical pathways through which such signals could reach S1. Several observations argue against an explanation based solely on trivial visual responses or peripheral reafference. First, many action observation-responsive neurons showed activity patterns that were congruent with execution-related responses in epochs lacking salient visual events (e.g., the Item Hold End epoch, when the hand was stationary on the object, as shown in Figure 4h and 5h), arguing against a purely stimulus driven visual response. Second, EMG recordings confirmed the absence of detectable covert muscle activity during observation trials, indicating that the modulation was not attributable to subtle movement-related peripheral reafference. Together, these findings argue against both visual and peripheral reafference-driven explanations, leaving cortico-cortical motor- or sensorimotor-related influences as more likely sources. Previous human studies have reported that tactile regions in S1 can be modulated by visual stimuli related to touch or body-related sensory events (Hansson et al. (2009); Kuehn et al. (2018)), and that somatosensory responses can be influenced by visual context (Rosenthal et al. (2023)). These findings are consistent with the possibility that visual information can influence S1 activity. However, based on the congruence between execution- and observation-related activity observed in the present study, it is unlikely that the modulation observed here is explained by visual input alone. Although the precise origin and content of the action observation-related signals cannot be unequivocally determined from the present data, the findings suggest that S1 activity during action observation reflects broader sensorimotor network dynamics, rather than functioning solely as a passive recipient of ascending somatosensory input.

### 3.2 A hierarchical gradient of action observation-related activity across S1

By examining different subdivisions across S1, we found that action observation-responsive neurons were more prevalent in areas 1 and 2 than in areas 3a and 3b. This difference may reflect the hierarchical organization of somatosensory processing within S1. Areas 3a and 3b are generally regarded as earlier stages of cortical processing for proprioceptive and cutaneous signals, respectively, whereas areas 1 and 2 are associated with progressively more integrative processing. Previous studies have shown that neurons in areas 1 and 2 exhibit broader receptive fields (Iwamura et al. (1983, 1985, 1993)) and more complex response properties, including mixed responses to cutaneous and proprioceptive stimuli (Hyvärinen and Poranen (1978); Costanzo and Gardner (1980); Warren et al. (1987); Kim et al. (2015)). Such properties likely reflect increasing integration of information across modalities and body segments along the somatosensory hierarchy. The greater proportion of action observation-responsive neurons in areas 1 and 2 therefore suggests that action observation-related signals may preferentially influence later stages of somatosensory processing, where cortical integration is more prominent. One possibility is that these higher-order subdivisions are more susceptible to cortico-cortical influences arising from motor and parietal areas during action observation. In contrast, the more strongly afferent-driven circuitry of areas 3a and 3b may limit the extent to which such modulation is expressed. Overall, the hierarchical gradient observed in the present study suggests that action observation-related activity is not uniformly distributed across S1, but instead depends on the functional organization and local circuitry of individual subdivisions.

### 3.3 Congruence of neuronal activity during action execution and observation

In premotor areas, neurons responsive during action observation often exhibit congruent activity during both action execution and observation (Gallese et al. (1996); Rizzolatti et al. (1996)). Such congruence may provide insight into the nature of the signals reaching S1 during action observation. In the present study, many neurons responsive during action observation exhibited activity patterns that were partially congruent between execution and observation conditions, although the magnitude and consistency of modulation were generally weaker during observation. This similarity to previously described premotor responses raises the possibility that motor-related signals distributed through the broader sensorimotor network contribute to the observed S1 activity. At the same time, substantial differences across S1 subdivisions were also observed. In particular, neurons in area 2 showed greater incongruence between execution and observation conditions compared with other subdivisions. The greater execution-observation incongruence suggests that area 2 may engage in sensorimotor processing distinct from that of other S1 subdivisions and may receive a broader range of cortico-cortical influences during action observation. Given that area 2 receives dense inputs from both motor cortex (Gharbawie et al. (2011)) and parietal cortex (Padberg et al. (2019)), the observed modulation may reflect more integrated sensorimotor influences arising through cortico-cortical interactions. Further studies will be necessary to clarify how these signals are generated and how they relate to the functional roles of different S1 subdivisions during action observation.

### 3.4 Functional implications of action observation-related activity in S1

The present study aimed to demonstrate the presence of neurons that respond during action observation in S1. Previous discussions of action observation-related activity have often focused on motor and cognitive processes associated with premotor and parietal cortices, including action understanding, motor representation, and social cognition (Jeannerod (2001); Brass and Heyes (2005); Rizzolatti and Craighero (2004)). Compared with these higher-order association regions, S1 is more directly involved in somatosensory processing, situated at the interface between peripheral sensory input and motor-related cortical computations. Therefore, the most straightforward interpretation of the present findings is that action observation-related activity in S1 reflects sensorimotor processes associated with observing biologically relevant actions. Such activity may reflect the engagement of distributed sensorimotor networks during action observation, including interactions between somatosensory, motor, and visuomotor cortical areas. Although the precise functional significance of these responses remains unresolved, the present observations demonstrate that action observation-related activity extends into early somatosensory cortex at the single-neuron level, broadening the range of cortical regions known to participate in action observation.

## 4 Methods

### 4.1 Experimental procedures

All experimental procedures were approved by the local Animal Research Committee at National Center of Neurology and Psychiatry, Japan, and carried out in accordance with Guidelines for Animal Experimentation in Neuroscience, which is compliant with The Act on Welfare and Management of Animals, and The Standards Relating to the Care and Management of Laboratory Animals and Relief of Pain of Japan (cf, https://www.jnss.org/en/animal_guidelines?u=e133dd8571fb8b88ab2b3fcd0b43dfed). Experiments involved two adult purpose-bred Japanese monkeys (macaca fuscata) (monkey Mu, male 9.5kg; monkey Ab, male 8.0 kg).

### 4.2 Behavioral task

The task involved an execution trial in which the monkey grasped different objects, and an observation trial in which the monkey observed the performance of the grasp by a human experimenter. We used the TEMPO system (Reflective Computing, WA, USA) to control a custom-made behavioral carousel and record behavioral events. The monkey and human experimenter were seated, facing each other across a custom-built carousel device. The carousel rotated to move an object on either the monkey or human side; in execution trials the object was presented to the monkey, whereas in observation trials the object was presented to the human experimenter (Figure 1A). We presented two different objects to each animal; ring and precision grip objects (Figure 1B) for monkey Mu, and ball-shaped and precision grip objects (Figure 1B) for monkey Ab. We did not aim to find a consistent neuronal activity for the same object in both animals, but rather to examine whether the neuronal activity could be changed for different objects for execution and observation conditions. As such, because each monkey preferred different types of grasp objects, we took advantage of the preference to facilitate the behavioral training.

In the execution condition, the trial began when both the monkey and the human experimenter placed one hand on the homepad for a certain period of time (100 ms). After a delay period (1000 ms), the object was rotated to the front of the monkey and turned visible through an opaque screen switched to be electronically transparent, with a LED illumination around the object (Object Presentation). After another period of delay (1000 ms), the LED color changed and a beep sound was presented (Go Cue), which instructed the monkey to reach out, grasp and pull the object. The onset of reach was defined as the time the monkey released the homepad (Reach Onset). The Middle of Reach was defined as the time in the middle of Reach Onset and the time the monkey touched the object (Item Touch), detected by a custom-made electrostatic sensor. The monkey held the object steadily for between 500 and 1000 ms until the next cue was given. The Object Hold (Item Hold) was defined as the time when the monkey pulled the object to the 90% of range of motion, which was measured by a distance sensor (E32-DC200F4R, E3X-HD6, OMRON). On the beep sound and LEDs turning off, the monkey released the object (Release Cue), returning the right hand on the homepad. The end of hold was defined as the time the lever of the object returned below 10% of range of motion (Item Hold End) based on the distance sensor. During the whole sequence of the trial, the experimenter was required to keep his/her right hand on the homepad. On successful completion of the sequence of the trial, the monkey was rewarded with diluted apple sauce. In the observation condition, the roles of the monkey and the human experimenter were reversed. As such, if the monkey moved its hand away from the homepad, the trial was considered a failure, with no reward being given.

### 4.3 Surgical procedures and electrophysiological recordings and stimulation

Extracellular single-unit recordings were made in the hand region of left S1 using a standard technique (Oya et al. (2020)). Briefly, under aseptic conditions, we made craniotomy over the pre/post central gyri of the left hemisphere to attach a recording chamber over the hand regions of S1 with dental acrylic. After the animal recovered, on a separate day we performed another surgery to implant cuff electrodes around the peripheral nerve branches: 1) superficial branch (SR), 2) deep branch (DR), and 3) the more proximal stem of the radial nerve (R), as well as pairs of wire electrodes (AS631, Cooner wire, CA, USA) into forearm muscles (Extensor Digitorum Communis (EDC), red: Extensor Carpi Ulnaris (ECU), magenta: Flexor Carpi Radialis (FCR), cyan: Flexor Digitorum Superficialis (FDS), and green: Triceps Brachii (Triceps)) for electromyographic (EMG) recordings. We first performed mapping sessions to identify target regions within S1, followed by neuronal recording sessions. For the mapping and recording sessions, we restricted head movement by securing the monkey’s head with a thermoplastic mask (Uni-frame, Toyo Medic, Tokyo, Japan) adjusted to fit each monkey (Drucker et al. (2015)). A hydraulic micromanipulator (MO-973; NARISHIGE, Japan) was attached to the recording chamber via a custom-made X-Y positioning stage. We performed mapping sessions to identify the distal hand region of S1 using SR and DR nerve stimulation (0.5-ms bipolar pulse), as well as tactile exploration by manual palpation. On each recording day, a tungsten microelectrode (Alpha Omega Engineering, Ltd, Israel) was advanced through S1 while the monkey was performing the task, and neurophysiological recordings were made using Alpha-Omega SnR System (Alpha Omega Engineering, Ltd, Israel), with an Ag-AgCl ball electrode being placed on the scar tissue grown over the cortical surface. The neural signal was amplified (x20) and digitized at 44 kHz for the neuronal signal. The EMG signals were concurrently recorded at a sampling rate of 5500 Hz with a band-pass filter between 20-2750 Hz. During the task, neuronal units were isolated by manually setting spike-detection thresholds, and thereafter examined as to whether they responded to electrical stimulation of the SR or DR nerve. The isolated units were sorted offline by isolating clusters within a space defined by the top three principal components extracted from the spike waveforms (Offline Sorter ver 2; Plexon, USA). Well isolated units were analysed further as datasets.

### 4.4 Classification of S1 subdivisions

Areas 3b, 1, and 2 were distinguished based on electrode depth, receptive field progression, and response properties, following Bensmaia et al. (2008) and Pei et al. (2010). As electrodes descended from the cortical surface, receptive fields shifted systematically from distal finger pads toward more proximal pads and palmar representations as electrodes advanced from area 1 into area 3b, while within 3b receptive fields progressed back from proximal, medial, and distal pads. The dissociation of areas 1 and 3b is corroborated by the SEPs evoked by stimulation of SR; generally the SEPs in area 1 have a positive deflection before a negative deflection, whereas ones in 3b have a negative deflection without the positive deflection in advance (Figure 1D). This change in SEP waveform was typically observed at approximately the same depth where the somatotopic progression of receptive fields reversed between areas 1 and 3b. The close correspondence between these two landmarks allowed us to use SEP waveform characteristics to identify the cortical area during behavioral recording sessions. Area 2 was identified more posterior-laterally, where neurons exhibited larger receptive fields and multimodal responses to both cutaneous and joint stimulation. The delineation described above was confirmed with post mortem histological analysis on monkey Mu.

### 4.5 Data analysis

The isolated neuronal firing of each identified unit was evaluated by constructing peri-event time histograms (PETHs) aligned to the eight behavioral events defined above, using a temporal window of 2000 ms (−1000 to +1000 ms relative to each event) and a bin size of 20 ms. Neuronal modulation was assessed in a 200-ms window (−100 to +100 ms relative to the event). Statistical significance was determined by estimating the probability that two consecutive bins exceeded the confidence interval defined by the baseline firing rate (calculated from −200 to −100 ms relative to object presentation). The confidence interval was set using the standard deviation (SD) of baseline activity (e.g., 2 SD approximately corresponding to an *α* level of 0.05).

The probability of observing such events by chance was estimated using the binomial distribution:

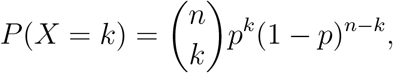

where n is the number of bins and p is the probability of a bin exceeding the threshold. For n = 10 bins (200-ms window) and p = 0.05 (2 SD), the probability of observing exactly two bins exceeding the threshold was calculated using the binomial probability density function in MATLAB: binopdf(2,10,0.05) = 0.187 (Baker et al., 2003). Of the 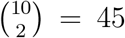 possible bin pairs, only 9 represented adjacent bins, yielding a proportion of 9*/*45 = 0.2. Thus, the probability of observing two consecutive supra-threshold bins by chance was estimated as 0.187 *×* 0.2 = 0.037. The same procedure was applied for other confidence levels (e.g., 2.33 SD for *α*=0.02, 2.58 SD for *α*=0.01, 2.81 SD for *α*=0.005).

To control for false positives arising from multiple comparisons across the seven different epochs, we corrected probabilities using the false discovery rate (FDR). FDR correction was performed according to the procedures described by Benjamini and Hochberg (1995) and Benjamini and Yekutieli (2001), using a q-value of 0.065.

Only neurons with a sufficient number of trials (15 trials) in all four categories, i.e., two grasp objects (ring and precision grip for monkey Mu; power grip and precision grip for monkey Ab) and two task conditions (Execution and Action Observation), were retained for analysis. This criterion yielded 195 neurons from monkey Mu and 179 neurons from monkey Ab. We further restricted the dataset to neurons that exhibited significant modulation in at least one epoch during execution of the grasping task relative to baseline activity (200 ms before object presentation), resulting in 190 neurons from monkey Mu and 170 neurons from monkey Ab. For each neuron, significant modulation in at least one of the seven defined epochs for a given grasp object under either Execution or Action Observation resulted in classification as modulated in that category. Neurons exhibiting significant modulation during both execution and action observation were further classified as action observation-responsive.

### 4.6 Exclusion of putative covert muscle activation with EMG analysis

To examine whether neuronal firing during action observation was influenced by covert movements that could explain the observed modulation, we analyzed the EMG activity during the same periods as those used for neuronal firing analysis. EMG signals were downsampled to 5,000 Hz and subjected to the same statistical analyses used for neuronal firing activity. Because EMG signals are continuous data, we evaluated whether activity from any recorded muscle exceeded the 2 SD confidence interval within the same time windows used for neuronal firing across the seven epochs, applying the same criterion for FDR correction. If a neuron was recorded in a session where EMG activity in any concurrently recorded muscle exceeded the threshold, that neuron was excluded from further analysis.

## Supplementary information

**Fig. S1.**
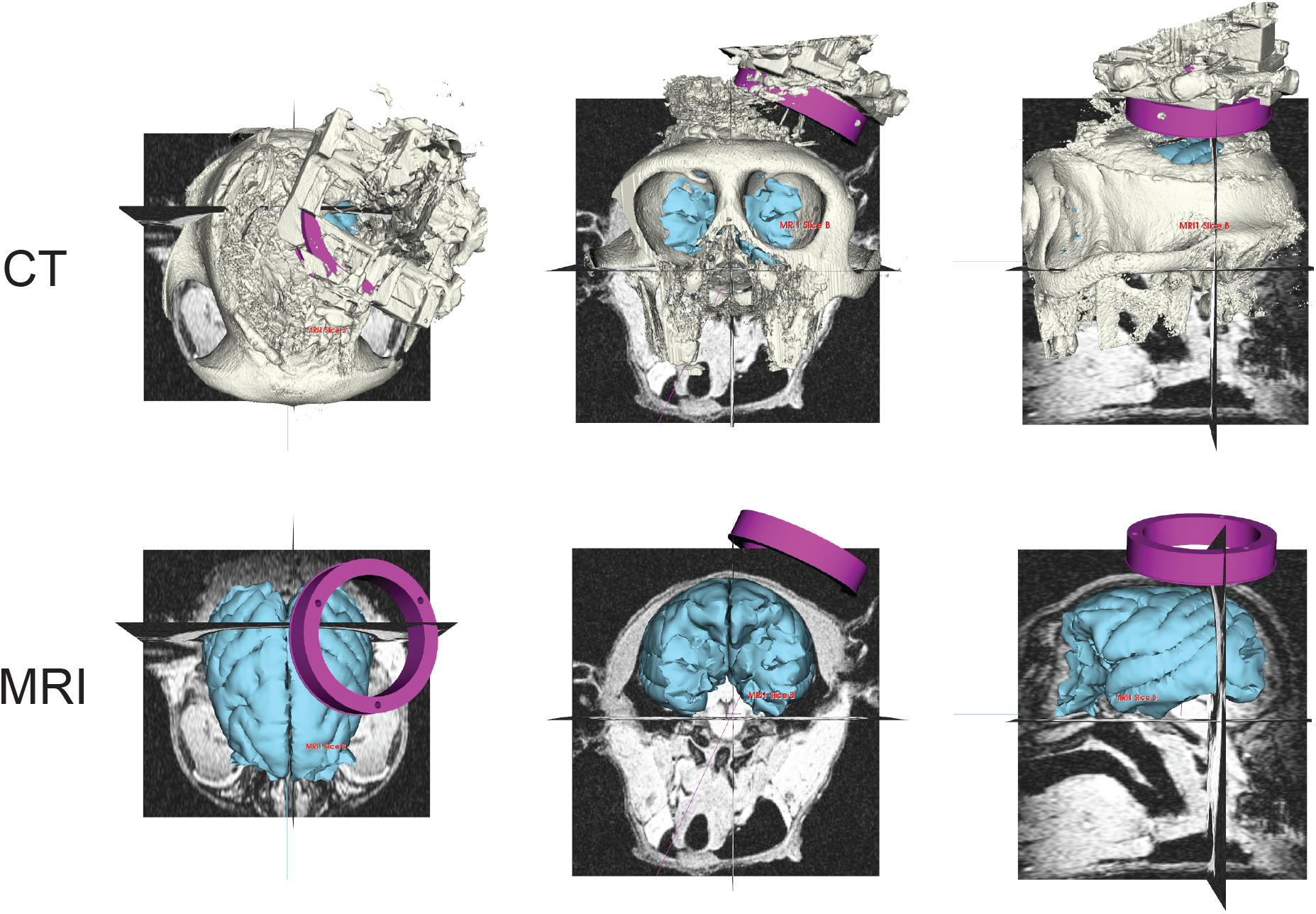
Projection of electrode insertion into S1 based on co-registration of CT and MRI.

**Fig. S2.**
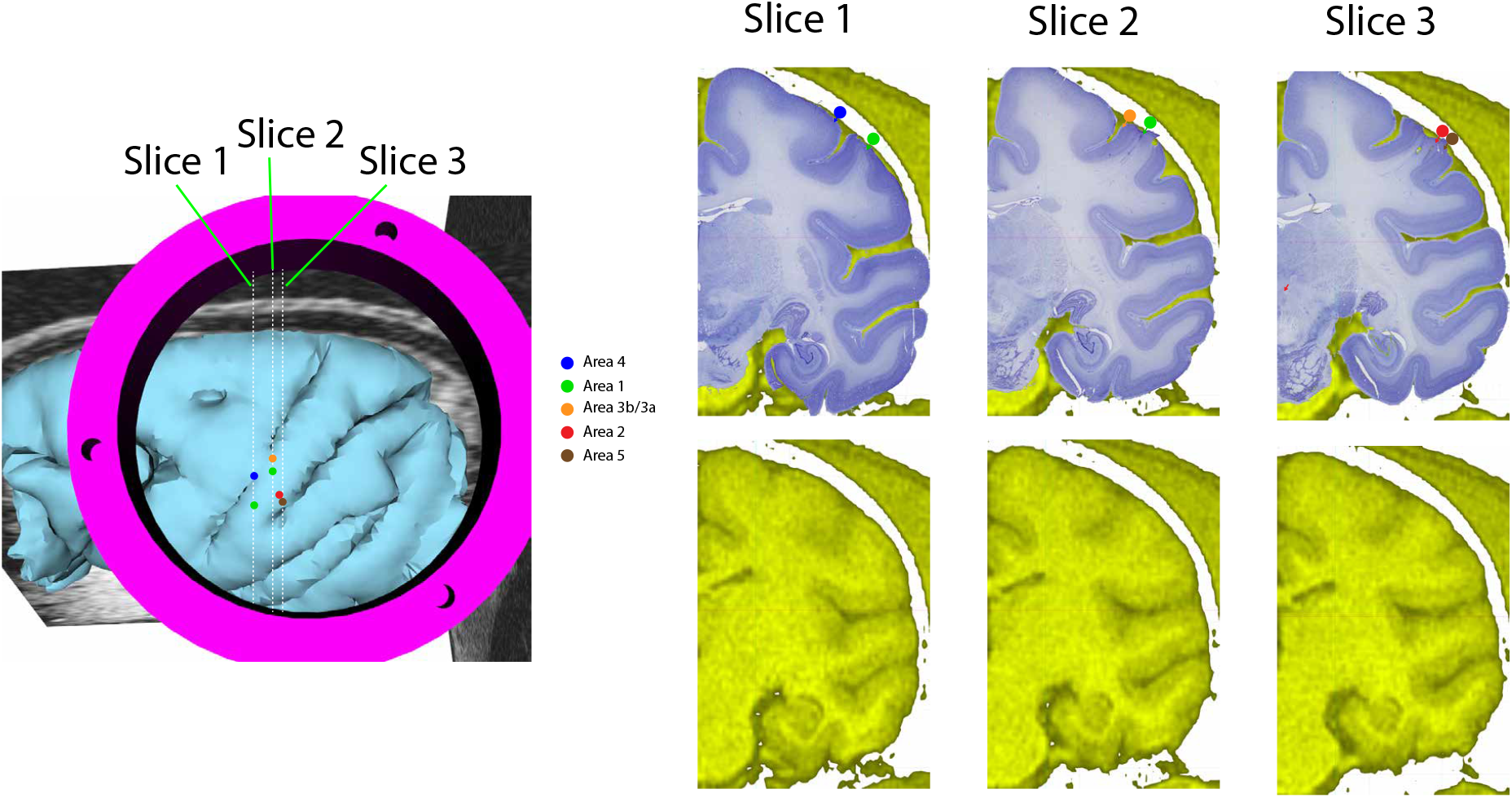
Confirmation of recorded areas by co-registration of MRI reconstruction and histology. The colored points stand for the places where electrolytic lesions were made. The whole brain slices were obtained from monkey Mu through formalin perfusion, and stained with Nissl staining technique.

## Acknowledgements

We are grateful to Chika Sasaki and Moweko Kudo for their technical support and assistance with experiments. This study is supported by a grant-in-aid from the Japan Society for the Promotion of Science (JSPS) (grant numbers 25K24607, 24K21313, 19K21825, 19H05724, 19H01092, 26120003 [to K. S.]).

## Contribution

T.O. contributed to experimental design, performed experiments, analyzed the data, prepared the figures, and wrote the original draft. A.Y. contributed to data acquisition and analysis. J.C. conceptualized the study, designed the experiments, and contributed to data acquisition and analysis. Sh.Ku. contributed to data acquisition and reviewed and edited the manuscript. Sa.Ki. contributed to data acquisition. K.S. conceptualized and supervised the project and edited the manuscript. All authors reviewed and approved the final manuscript.

## Notes

### Competing Interest Statement

The authors have declared no competing interest.

